# Lateral gene transfer drives metabolic flexibility in the anaerobic methane oxidising archaeal family *Methanoperedenaceae*

**DOI:** 10.1101/2020.02.06.936641

**Authors:** Andy O. Leu, Simon J. McIlroy, Jun Ye, Donovan H. Parks, Victoria J. Orphan, Gene W. Tyson

## Abstract

Anaerobic oxidation of methane (AOM) is an important biological process responsible for controlling the flux of methane into the atmosphere. Members of the archaeal family *Methanoperedenaceae* (formerly ANME-2d) have been demonstrated to couple AOM to the reduction of nitrate, iron, and manganese. Here, comparative genomic analysis of 16 *Methanoperedenaceace* metagenome-assembled genomes (MAGs), recovered from diverse environments, revealed novel respiratory strategies acquired through lateral gene transfer (LGT) events from diverse archaea and bacteria. Comprehensive phylogenetic analyses suggests that LGT has allowed members of the *Methanoperedenaceae* to acquire genes for the oxidation of hydrogen and formate, and the reduction of arsenate, selenate and elemental sulfur. Numerous membrane-bound multi-heme *c* type cytochrome complexes also appear to have been laterally acquired, which may be involved in the direct transfer of electrons to metal oxides, humics and syntrophic partners.

**Importance:** AOM by microorganisms limits the atmospheric release of the potent greenhouse gas methane and has consequent importance to the global carbon cycle and climate change modelling. While the oxidation of methane coupled to sulphate by consortia of anaerobic methanotrophic (ANME) archaea and bacteria is well documented, several other potential electron acceptors have also been reported to support AOM. In this study we identify a number of novel respiratory strategies that appear to have been laterally acquired by members of the *Methanoperedenaceae* as they are absent in related archaea and other ANME lineages. Expanding the known metabolic potential for members of the *Methanoperedenaceae* provides important insight into their ecology and suggests their role in linking methane oxidation to several global biogeochemical cycles.

## Introduction

Anaerobic oxidation of methane (AOM) is an important microbiological process moderating the release of methane from anoxic waters and sediments into the atmosphere (1–4). Several diverse uncultured microbial lineages have been demonstrated to facilitate AOM. The bacterium “*Candidatus* Methylomirabilis oxyfera” is proposed to couple AOM to denitrification from nitrite, generating oxygen from nitric oxide for the activation of methane (5). Different lineages of anaerobic methanotrophic (ANME) archaea are hypothesised to mediate AOM through the reversal of the methanogenesis pathway and conserve energy using mechanisms similar to those found in methylotrophic and aceticlastic methanogens (6). Unlike methanogens, most of these ANMEs encode a large repertoire of multi-heme *c*-type cytochromes (MHCs), which are proposed to mediate direct interspecies electron transfer to syntrophic sulfate-reducing bacteria (SRB)(7, 8), and/or the reduction of metal oxides and humic acids (9–12).

Currently, several clades within the archaeal phylum Euryarchaeota have been shown to be capable of anaerobic methanotrophy and include ANME-1a-b, ANME-2a-c, *Methanoperedenaceae* (formerly known as ANME-2d), and ANME-3 (refs. 13, 14, 15). Marine ANME lineages are often observed to form consortia with SRBs, with ANME-1 and ANME-2 (a,b, and c) being associated with multiple genera within Desulfobacterales and *Desulfobulbaceae* (13, 16–20), thermophilic ANME-1 with “*Candidatus* Desulfofervidus auxilii” (8, 21) and ANME-3 with SRBs of the *Desulfobulbus* (22). While members of the family *Methanoperedenaceae* have also recently been associated with SRB of the family *Desulfobulbaceae* in a freshwater lake sediment (23), they also appear to oxidise methane independently using a range of electron acceptors. The type species of this family, “*Candidatus* Methanoperedens nitroreducens”, was originally enriched in a bioreactor and shown to couple AOM to the reduction of nitrate via a laterally transferred nitrate reductase (15). Subsequently, “*Ca*. Methanoperedens sp. BLZ1” was also found to encode a laterally transferred nitrite reductase, which is also present in the genome of “*Ca*. M nitroreducens”, potentially allowing these microorganisms to coupled AOM to dissimilatory nitrate reduction to ammonia (DNRA) (24). More recently, three novel species belonging to the *Methanoperedenaceae* were enriched in bioreactors demonstrated to couple AOM to the reduction of insoluble iron or manganese oxides (9, 12). These microorganisms did not encode dissimilatory nitrate reduction pathways, but instead were inferred to use multiple unique MHCs during metal-dependent AOM to facilitate the transfer of electrons to the metal oxides (9, 12), consistent with the extracellular electron transfer mechanisms proposed for marine ANME (7, 8). Bioreactor performance and 16S rRNA gene amplicon data has also been used to suggest that members of the *Methanoperedenaceae* are capable of AOM coupled to the reduction of selenate and chromium(VI), although this remains to be confirmed with more direct evidence (25, 26). Notably, members of the *Methanoperedenaceae* have been observed to facilitate AOM coupled to multiple terminal electron acceptors within the same natural sediment (27). Individual members of the family can possess such metabolic flexibility, with a lab-enriched species shown to couple AOM to the reduction of nitrate, iron and manganese oxides (10). Given the relatively poor genomic representation of the *Methanoperedenaceae*, and the lack of detailed physiological studies of its members, it is likely that considerable metabolic diversity for the lineage remains to be discovered.

In this study, comparative analysis was conducted on 16 *Methanoperedenaceae* metagenome-assembled genomes (MAGs) recovered from various environments to investigate the metabolic diversity and versatility of the family and to understand the evolutionary mechanisms responsible for these adaptations. These analyses indicate that members of the *Methanoperedenaceae* have acquired a large number of genes through LGT that potentially allow AOM to be coupled to a wide range of electron acceptors, suggesting their role in methane oxidation extends beyond environments with nitrate and metal oxides.

## Results and Discussion

### Expanding the genomic representation of the Methanoperedenaceae

In order to explore the metabolic diversity within the *Methanoperedenaceae*, comparative genomic analysis was performed on both publicly available and newly acquired MAGs (**Table 1**). The publicly available genomes include five MAGs recovered from bioreactors where AOM is coupled to the reduction of nitrate (“*Ca*. Methanoperedens nitroreducens”; M.Nitro (15), and “*Ca*. Methanoperedens sp. BLZ2”; BLZ2 (ref. 28)), iron (“*Ca*. Methanoperedens ferrireducens”; M.Ferri (9)) and manganese (“*Ca*. Methanoperedens manganicus” and “*Ca*. Methanoperedens manganireducens”, Mn-1 and Mn-2, respectively (12)). Also included are two environmental MAGs recovered from groundwater samples from the Horonobe and Mizunami underground research laboratories in Japan (HGW-1 and MGW-1) (29, 30), and one MAG from an Italian paddy soil sample (IPS-1) (31). In order to recover additional genomes belonging to the family, GraftM (32) was used to screen public metagenome sequence datasets from NCBI for *Methanoperedenaceae*-related 16S rRNA and *mcrA* gene sequences. Subsequent assembly and genome binning on datasets found to contain *Methanoperedenaceae*-like sequences led to the recovery of an additional eight MAGs belonging to the family. Six of these were from arsenic contaminated groundwater samples (ASW-1-6), and a further two from sediment and groundwater samples from a copper mine tailings dam (CMD-1 and CMD-2). All 16 MAGs are highly complete (≥87.4%) with low contamination (≤5.9%) based on 228 Euryarchaeota-specific marker genes (**Table 1**)(33). These genomes vary in GC content from 40.2 to 50.7% and range in size from 1.45 to 3.74 Mbp.

**Table 1.** Characteristics of the metagenome-assembled genomes.

| Bin Id | Genome size (Mbp) | No. scaffolds | N50 (scaffolds; bp) | Strain heterogeneity<sup>#</sup> | Compl. (%)<sup>#</sup> | Cont. (%)<sup>#</sup> | GC | #CDS | Environment | Accession no.* | 16S rRNA gene? |
| --- | --- | --- | --- | --- | --- | --- | --- | --- | --- | --- | --- |
| ASW-1 | 1.52 | 271 | 7,386 | 0.0 | 87.5 | 0.0 | 47.8 | 1946 | Arsenic contaminated groundwater (Bangladesh) | SRR1563167, SRR1564103, SRR1573565, SRR1573578, SAMN10961276 | N |
| ASW-2 | 2.63 | 157 | 28,058 | 25.0 | 94.4 | 4.8 | 48.0 | 2944 | Arsenic contaminated groundwater (Bangladesh) | SRR1563167, SRR1564103, SRR1573565, SRR1573578, SAMN10961277 | N |
| ASW-3 | 2.51 | 100 | 44,967 | 0.0 | 100.0 | 1.3 | 50.7 | 2892 | Arsenic contaminated groundwater (Bangladesh) | SRR1563167, SRR1564103, SRR1573565, SRR1573578, SAMN10961278 | N |
| ASW-4 | 2.24 | 155 | 24,336 | 0.0 | 97.1 | 0.7 | 43.2 | 2464 | Arsenic contaminated groundwater (Bangladesh) | SRR1563167, SRR1564103, SRR1573565, SRR1573578, SAMN10961279 | N |
| ASW-5 | 2.97 | 221 | 19,046 | 0.0 | 95.0 | 2.6 | 48.9 | 3353 | Arsenic contaminated groundwater (Bangladesh) | SRR1563167, SRR1564103, SRR1573565, SRR1573578, SAMN10961280 | N |
| ASW-6 | 2.19 | 68 | 56,691 | 66.7 | 99.4 | 2.0 | 46.6 | 2472 | Arsenic contaminated groundwater (Bangladesh) | SRR1563167, SRR1564103, SRR1573565, SRR1573578, SAMN10961281 | Y |
| BLZ1** | 3.74 | 514 | 17,508 | 13.33 | 96.73 | 6.56 | 40.2 | 4659 | AOM-nitrate reactor (Netherlands) | LKCM00000000.1 | Y |
| BLZ2 | 3.74 | 85 | 74,304 | 0.0 | 99.4 | 4.6 | 40.3 | 4041 | AOM-nitrate reactor (Netherlands) | GCA_002487355.1 | N |
| CMD-1 | 1.85 | 116 | 27,949 | 100.0 | 98.0 | 0.7 | 44.9 | 2261 | Copper mine tailings dam (Brazil) | SRR5161805, SRR5161795, SAMN10961282 | N |
| CMD-2 | 1.45 | 221 | 9,704 | 0.0 | 88.4 | 0.0 | 44.1 | 1786 | Copper mine tailings dam (Brazil) | SRR5161805, SRR5161795, SAMN10961283 | N |
| HGW-1 | 2.00 | 128 | 24,496 | 33.3 | 96.4 | 2.0 | 43.2 | 2288 | Groundwater samples (Japan) | GCA_002839545.1 | Y |
| IPS-1 | 3.52 | 250 | 27,331 | 10.0 | 97.7 | 5.9 | 44.1 | 3970 | Paddy field soil (Italy) | GCA_900196725.1 | Y |
| M.Ferri | 2.91 | 59 | 88,069 | 0.0 | 98.7 | 1.3 | 40.8 | 3019 | AOM-iron reactor (Australia) | GCA_003104905.1 | Y |
| M.Nitro | 3.20 | 10 | 54,4976 | 0.0 | 99.7 | 1.3 | 43.2 | 3428 | AOM-nitrate reactor (Australia) | GCA_000685155.1 | Y |
| MGW-1 | 2.08 | 161 | 17,186 | 0.0 | 97.4 | 3.6 | 44.8 | 2488 | Groundwater samples (Japan) | Not available<sup>§</sup> | N |
| Mn-1 | 3.59 | 68 | 87,551 | 0.0 | 100.0 | 1.3 | 40.6 | 3737 | AOM-manganese reactor* (Australia) | SAMN10872768 | N |
| Mn-2 | 3.32 | 116 | 49,809 | 0.0 | 99.4 | 4.6 | 42.9 | 3684 | AOM-manganese reactor* (Australia) | SAMN10872769 | N |

<sup>#</sup>Completeness (compl.), contamination (cont.), and strain heterogeneity were estimated using CheckM (33)
<sup>\*</sup>Genome accession numbers. For the MAGs assembled in this study the SRA accession numbers are also given.
<sup>\*\*</sup>The BLZ1 genome was not used in analyses as it is almost identical to the BLZ2 genome (99.5% ANI) and has inferior completeness and contamination values. The BLZ1
bioreactor was the parent system of the BLZ2 bioreactor.
<sup>§</sup>This genome was provided by Dr Yohey Suzuki and is associated with the study of Hernsdorf and colleagues (29)

A genome tree including 1,199 publicly available archaeal genomes, based on a concatenated set of 122 marker genes (34), confirmed the phylogenetic placement of the 16 MAGs within the *Methanoperedenacae*. The genome tree supports that these MAGs form a monophyletic clade sister to the GoM-Arc1 genomes (**Figure 1**). These genomes likely represent three separate genera within the family, based on their placement within a reference tree, relative evolutionary distance, FastANI distance, and average amino acid identity (AAI (35); 61.3 to 89.2%; **Figure S1**). All MAGs were classified as members of the genus “*Ca*. Methanoperedens”, except HGW-1 and ASW-3 which appear to represent independent genus level lineages (**Figure 1**). Phylogenetic analysis of the six MAGs containing 16S rRNA genes was consistent with the genome tree (**Figure S2**), supporting their classification as members of the *Methanoperedenaceae* family.

**Figure 1.**
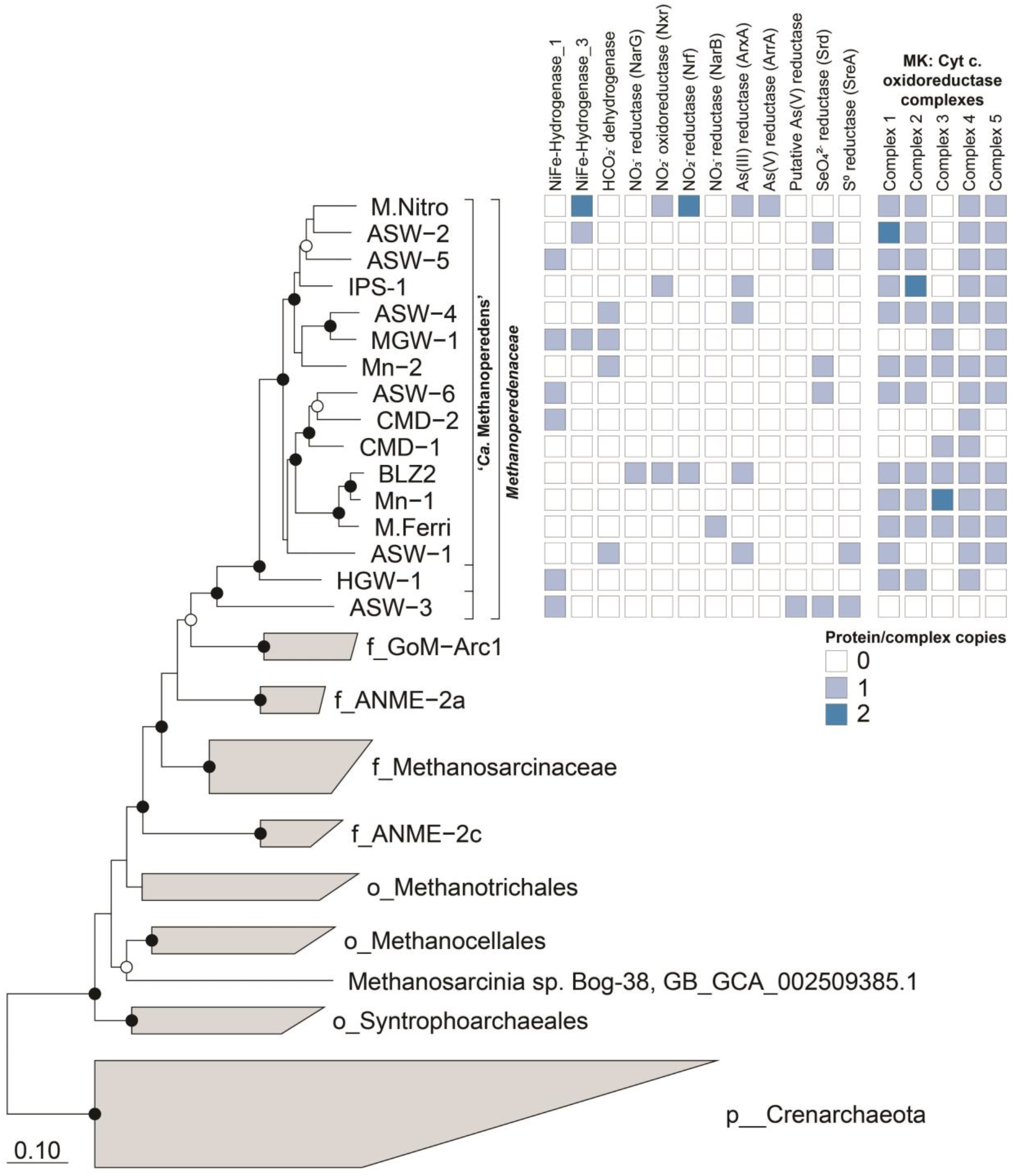
Phylogenetic placement of the *Methanoperedenaceae* MAGs and distribution of potential terminal electron acceptors. The genome tree was inferred using maximum-likelihood with a concatenated set of 122 archaeal specific marker genes. Black and white dots indicate >90% and >70% bootstrap values, respectively. The scale bar represents amino acids nucleotide changes. Based on GTDB-Tk the family *Methanoperedenaceae* includes three genera including “*Ca*. Methanoperedens” which are denoted with brackets. The table to the right of the tree shows the presence/absence of gene associated with potential terminal electron acceptors in each corresponding *Methanoperedenaceae* genome.

### Potential electron donors used by the Methanoperedenaceae

Metabolic reconstruction of the *Methanoperedenaceae* MAGs showed that all genomes encoded the central methanogenesis pathway, inclusive of the methyl-coenzyme M reductase, supporting their potential for the complete oxidation of methane to CO_2_ (**Figures 2 and S3**). The annotation of membrane-bound formate dehydrogenases (FdhAB) in five of the *Methanoperedenaceae* MAGs (Mn-2, ASW-4, ASW-1, MGW-1, and BGW-1; **Figure 3**) suggests that some members of the family may also oxidise formate (E_0_ [CO_2_/HCOO^-^] = – 430 mV) (36). As the enzyme is reversible, these species could also potentially produce formate as a supplementary electron sink during AOM. Formate was suggested as a putative electron shuttle between ANME-1 and their syntrophic partner SRB, based on the annotation and expression of an *fdhAB* in ANME-1, but this has not been supported with physiological studies (37, 38). The putative formate dehydrogenase encoded in the Mn-2 MAG is phylogenetically related to an FdhA found in the genome of *Caldiarchaeum subterraneum*, while those encoded by ASW-4, ASW-1, MGW-1, and BGW-1 appear to be more similar to the FdhA of *Methanocellaceae* archaeon UBA148 (**Figure 3**).

**Figure 2.**
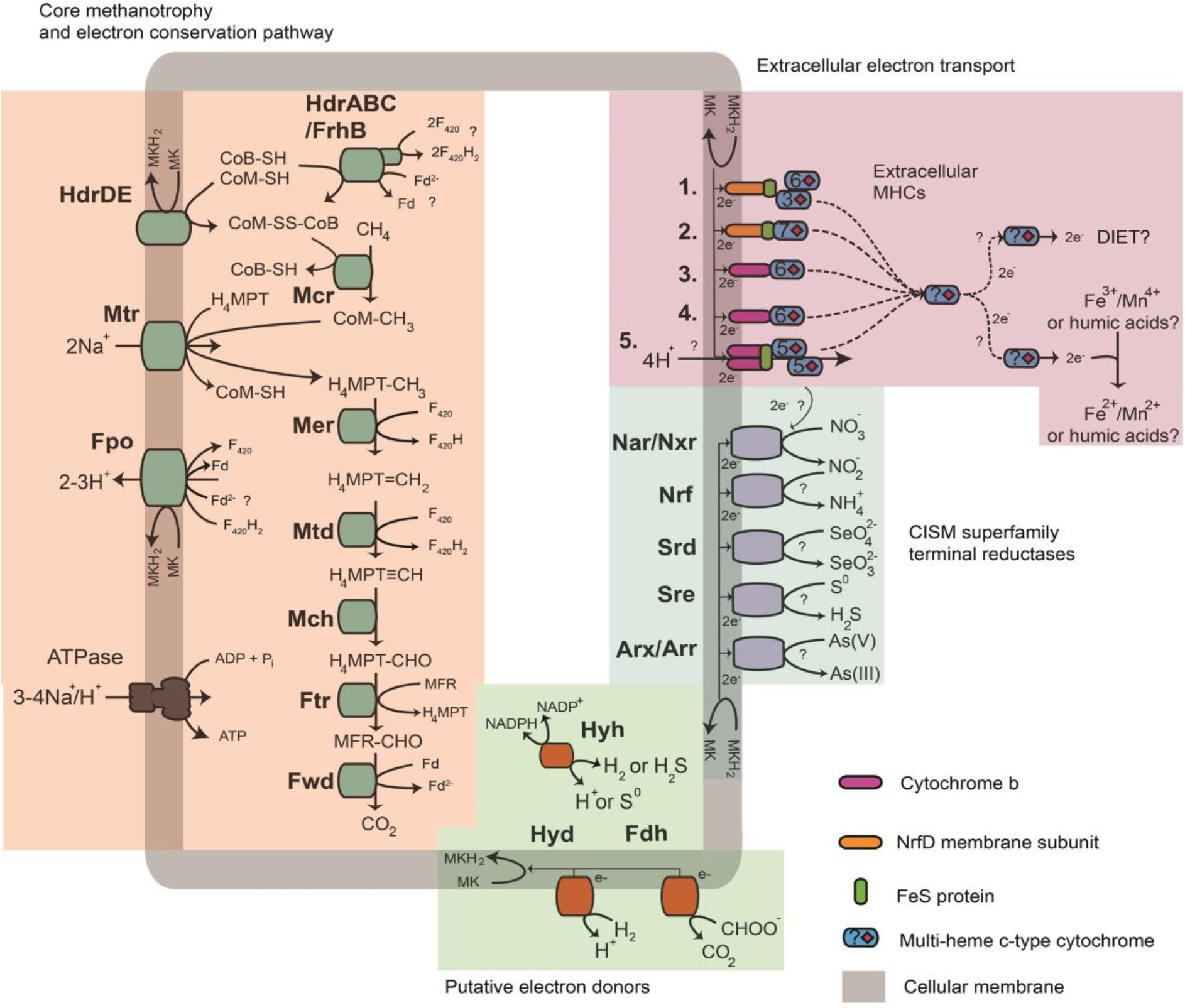
Metabolic capabilities of the *Methanoperedenaceae*. Key metabolic pathways for anaerobic oxidation of methane, energy conservation mechanisms, hydrogen and formate oxidation, and electron acceptors found within the pangenome of the *Methanoperedenaceae*. Numbers 1-5 indicate the different menaquinone:cytochrome c oxidoreductases conserved in the *Methanoperedeneceae* MAGs (**Dataset S1A**). Abbreviations for enzymes and co-factors in the figure are:H_4_MPT, tetrahydromethanopterin; MFR, methanofuran; Fwd, formyl-methanofuran dehydrogenase; Ftr, Formylmethanofuran/H_4_MPT formyltransferase; Mch, methenyl-H_4_MPT cyclohydrolase; Mtd, F_420_-dependent methylene H_4_MPT dehydrogenase; Mer, F_420_-dependent methylene-H_4_MPT reductase; Mtr, Na^+^-translocating methyl-H_4_MPT:coenzyme M methyltransferase; Mcr, methyl-coenzyme M reductase; F_420_, F_420_ coenzyme; Fd, ferredoxin; CoM-SH, coenzyme M; CoB-HS, coenzyme B; Hdr, heterodisulfide reductase; Fpo, F_420_H_2_ dehydrogenase; Hyd, type-1 NiFe hydrogenase; Hyh, type-3b NiFe hydrogenase; Fdh, formate dehydrogenase; Nar, nitrate reductase; Nrf, nitrite reductase, Ttr, tetrathionate reductase; Arx, arsenite oxidase; Arr, arsenate reductase; DIET, direct interspecies electron transfer.

**Figure 3.**
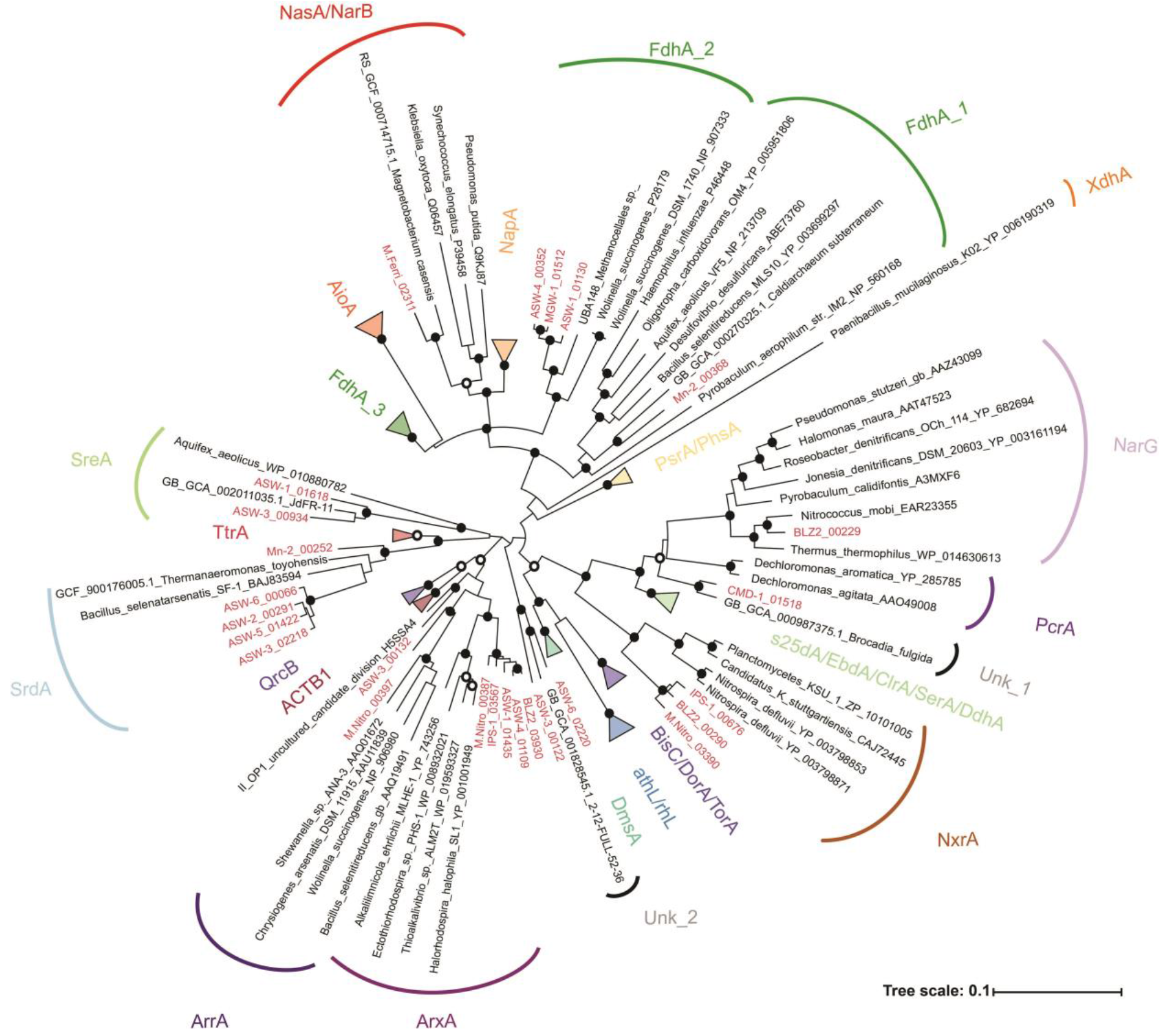
Phylogenetic analysis of the catalytic subunits of the CISM superfamily. Putative genes recovered from the *Methanoperedenaceae* are highlighted in red. The gene tree was inferred by maximum-likelihood and support values calculated via non-parametric bootstrapping. Black and white dots indicate >90% and >70% bootstrap support, respectively. The scale bar represents amino acid changes. ACTB1, alternate complex III, domain of subunit B; ArrA, arsenate reductase; ArxA, arsenite oxidase; AthL, Pyrogallol hydroxytransferase; BisC, biotin sulfoxide reductase; ClrA, chlorate reductase; EbdA, Ethylbenzene dehydrogenase; s25dA, C25dehydrogenase; DmsA, DMSO reductase; DorA, DMSO reductase; NapA, nitrate reductase; NarG, nitrate reductase; NasA, assimilatory nitrate reductase; NarB, assimilatory nitrate reductase; NxrA, nitrite oxidoreductase; PsrA, polysulfide reductase; PhsA, thiosulfate reductase; QrcB, quinone reductase complex; TtrA tetrathionate reductase; DmsA, PcrA, perchlorate reductase; SrdA, Selenate reductase; SreA, sulfreductase; TorA, TMAO reductase; XdhA, xanthine dhydrogenase; FdhA, formate dehydrogenase; rhL, Resorcinol hydroxylase; Unk, unknown putative reductase. Amino acid sequences are included in **Dataset S1B**.

The use of hydrogen (H_2_; E_0_ = −414mV (39)) as an electron source was previously suggested for MGW-1 and HGW-1 which encode Group 1 membrane-bound NiFe hydrogenase complexes, composed of a NiFe catalytic subunit, a FeS electron transfer subunit, and a membrane-bound *b*-type cytochrome (29, 30). These hydrogenases, along with similar Group 1 NiFe hydrogenases identified in the ASW-6 and CMD-2 MAGs, form a monophyletic clade with those encoded by the MAG for “*Ca*. Hydrothermarchaeota” (JdFR-18), which belongs to the archaeal phylum Hydrothermarchaeota (40), and several members of the Halobacterota (**Figure S4A**). The ASW-3 and ASW-5 MAGs encode Group 1 NiFe hydrogenases that are basal to Vho/Vht/Vhx hydrogenases encoded by members of the genus *Methanosarcina* (41). As the ASW-5 NiFe hydrogenase does not encode a *b*-type cytochrome (**Figure S4B**), it is unclear how electrons are derived from hydrogen. In addition to the membrane-bound NiFe hydrogenases, the M.Nitro MAG was found to encode genes for two different sets of Group 3b cytoplasmic hydrogenases (**Figure S4A**). The MGW-1 (ref. 29) and ASW-2 MAGs also encode Group 3b hydrogenases which have been implicated in hydrogen evolution and nicotinamide adenine dinucleotide phosphate (NADPH) reduction (42). Similar complexes have also been shown to have hydrogen oxidation and elemental sulfur reducing capabilities (42–44). It is unknown how these Group 3b hydrogenases would contribute to energy conservation given their predicted cytoplasmic localisation. The functionality of the annotated Group 1 and 3 NiFe hydrogenases is supported by the identification of the NiFe binding motifs (L1 and L2) on their NiFe catalytic subunits and the annotation of all or most of the hydrogenase maturation genes (hypA-F) on the same *Methanoperedenaceae* MAGs (**Dataset S1D**). The potential for some *Methanoperedenceae* to couple the oxidation of hydrogen and/or formate to the reduction of exogenous electron acceptors would be advantageous with the dynamic availability of methane in natural environments (45).

### Pathways for energy conservation during AOM in the Methanoperedenaceae

All members of the *Methanoperedenaceae* encode the Fpo complex (FpoABCDHIJ1J2LMNOF), a homolog of Complex I (nuoABCDEFGHIJKLMN), which is hypothesised to oxidize F420H2 coupled to the reduction of a membrane-bound soluble electron carrier, and translocation of two protons out of the cell (**Figures 2 and S5A**) (41, 46). While members of the *Methanosarcinales* and marine ANME-2a are reported to typically use methanophenazine (MP) as their membrane-bound soluble electron carrier, the *Methanoperedenaceae* and ANME-1 have previously been suggested to use menaquinone (MK) based on the annotation of the futalosine pathway for MK biosynthesis in several MAGs representing these lineages (47). Comparative genomic analysis of the 16 *Methanoperedenaceae* MAGs revealed that the futalosine pathway is a conserved feature of all members, except the most basal member ASW-3 (see later; **Dataset S1A**). As has previously been suggested by Arshad *et al*., (48), the larger difference in redox potential between F_420_ (*E_0_* = −360mV) and MK (*E_0_* = −80mV (49)), relative to F420 and MP (*E_0_* = - 165mV (50)), would theoretically allow the Fpo complex to translocate more protons (3H^+^/2e^-^) out of the cell for every molecule of F_420_ oxidised, giving a higher overall energetic yield from AOM (**Figure S5B**).

Phylogenetic analysis of the Fpo complex in the *Methanoperedenaceae* MAGs showed that the FpoKLMNO subunits are homologous to proteins found in MP utilising members of the *Methanosarcinales*. The FpoABCDHIJ1J2 subunits are more similar to those found in microorganisms known to use MK and other quinones, which have more positive redox potentials (**Figures S5 and S6**; **Dataset S1E**) (51). As the latter subunits (specifically FpoH) are responsible for interaction with the membrane soluble electron carrier pool (52, 53), this observation provides further support to the use of MK by members of the *Methanoperedenaceae*. To our knowledge, this is the first reported example of a lineage encoding a ‘hybrid’ Complex I homolog possessing subunits with homology to those found in phylogenetically diverse microorganisms (**Figure S6**). The GoM-Arc-I MAGs appear to possess the MK biosynthesis pathway and a similar ‘hybrid’ Fpo complex to the *Methanoperedenaceae* (**Figure S6**), suggesting that the evolutionary adaptation of the lineage to utilise MK occurred prior to the divergence of these two related families. Members of the GoM-Arc-1 clade possess Mcr-like complexes (**Figure S3**) and are suggested to use short-chain alkanes – possibly ethane (54, 55). Interestingly, the FpoMNO subunits of the ASW-3 MAG cluster with those of the other members of the *Methanoperedenaceae* family, while their FpoABCDHIJ1J2KL subunits are most similar to those of the ANME-2a and other members of the *Methanosarcinales* (**Figure S6**). While the genes involved in MP biosynthesis are not known, the absence of the MK biosynthesis pathway indicate that ASW-3 likely uses MP. As the most basal lineage of this family, ASW-3 may have adapted to use MP after the evolutionary divergence of the GoM-Arc-I and *Methanoperedenaceae*, although further genomic representation of this lineage is required to verify this hypothesis.

Comparative genomic analyses of the *Methanoperedenaceae* MAGs revealed that none of these genomes encode an Rnf complex, which is hypothesised to re-oxidise ferredoxin coupled to the transport of sodium ions out of the cell and the reduction of MP in marine ANME-2a (7, 56) and other methylotrophic methanogens (41, 57, 58). In the absence of this complex, ferredoxins could be re-oxidised with a ‘truncated’ Fpo complex, similar to the Fpo complex possessed by *Methanosaeta thermophila* (59). Alternatively an electron confurcating mechanism could be used for the re-oxidation of ferredoxin, coenzyme M, and coenzyme B, coupled to the reduction of two F420 via a cytoplasmic complex composed of a heterodisulfide reductase (HdrABC) and a F420 hydrogenase subunit B (FrhB) (24). The two additional F420H2 could subsequently be fed back into the Fpo complex, greatly increasing the overall bioenergetic yield (24) (**Figure 2**). All of the *Methanoperedenaceae* MAGs have the genetic potential for these alternate strategies for re-oxidation of ferredoxin during AOM, however, further experimental validation is required to test these hypotheses.

### Conservation of unique menaquinone: cytochrome c oxidoreductases within the Methanoperedenaceae

Five different putative MK:cytochrome c oxidoreductase gene clusters (**Figures 1 and 2**; **Dataset S1A**) that are hypothesised to mediate the transfer of electrons out of the cytoplasmic membrane were identified in the *Methanoperedenaceae* MAGs. These gene clusters include a non-canonical bc1/b6f complex adjacent to two hypothetical proteins and two 6-haem multi-heme cytochromes (MHCs; Group 1), two clusters where a *b-*type cytochrome is adjacent to a 6-haem MHC (Groups 2 and 3), and another two clusters where a NrfD-like transmembrane protein is adjacent to an electron transferring 4Fe-4S ferredoxin iron-sulfur protein and MHCs (Groups 4 and 5; **Figure 2**). These bc and NrfD complexes are frequently found in other metal reducing microorganisms and mediate electron transport from the cytoplasm to the periplasm (60–62).

Most of the 16 *Methanoperedenaceae* MAGs (except CMD-1 and ASW-3) have more than one of these MK:cytochrome oxidoreductase complexes and 11 have at least four (**Figure 1**). ASW-3 is the only MAG not to encode any MK: cytochrome *c* oxidoreductases, which is consistent with its putative use of MP. A gene encoding a cytochrome-*b* found to be most similar to “*Ca*. Methanohalarchaeum thermophilum” was identified in ASW-3; however, in the absence of a collocated MHC gene, the extracellular electron transfer step for this microorganism is unclear.

Phylogenetic analysis of the membrane-bound subunits of the MK:cytochrome *c* oxidoreductases (**Figure 2**), which include the NrfD subunits (from Groups 1 and 2) and the *b*-type cytochromes (from Groups 3, 4 and 5), showed that they have been potentially laterally transferred from diverse donors (**Figure S7**). The *Methanoperedenaceae* NrfD subunits formed independent clusters with sequences from members of the Dehalococcoidales family RBG-16-60-22 (Group 1) and a single MAG (RBG-16-55-9) from the candidate phylum Bipolaricaulota (Group 2; **Figure S7A**). The *b*-type cytochromes of the *Methanoperedenaceae* belong to three distinct clades (**Figure S7B**). The *b*-type cytochromes from Groups 3 and 4 clustered with proteins from GoM-ArcI, indicating vertical genetic inheritance from an ancestor of these two families, and Group 5 proteins clustered with those from the class Archaeoglobi (40).

The conservation of multiple conserved laterally transferred MK:cytochrome *c* oxidoreductases in most of the *Methanoperedenaceae* MAGs may contribute to the reported ability for members of the family to reduce a variety of electron acceptors with a range of redox potentials that include Fe(III) oxide reduction (−100mV to 100mV) (63), nitrate (+433mV)(24), and Mn(IV) (+380mV) (36). Transcriptomic analyses has shown that different MK:cytochrome *c* oxidoreductases are expressed in different species of the genus “*Ca*. Methanoperedens” during AOM coupled to the reduction of Fe(III) oxides (9), Mn(IV) oxides (12), and nitrate (15, 24). A similar phenomenon is observed for the species *Geobacter sulfurreducens*, where different extracellular electron pathways were used when reducing different electron acceptors (64).

### Potential electron acceptors used by the Methanoperedenaceae

Annotation of the *Methanoperedenaceae* MAGs revealed a wide array of genes associated with previously undescribed respiratory strategies for the family that appear to have been acquired via LGT. Principally, these are putative terminal oxidoreductase complexes belonging to the Complex-Iron-Sulfur-Molybdenum (CISM) superfamily that were absent in the genomes of related archaeal lineages (**Figure 3**). These complexes are composed of a catalytic subunit, an iron-sulfur protein, and a membrane-bound subunit, and facilitate the transfer of electrons between the electron acceptor/donor and the MK pool (**Figure 2**).

As previously reported, the MAGs M.Nitro, BLZ2, and IPS-1 encode respiratory nitrate reductases that are part of the CISM superfamily, allowing them to independently mediate AOM coupled to nitrate reduction (15, 24, 65). Based on phylogenetic analysis (**Figure 3**), genes encoding cytoplasmic nitrite oxidoreductases (NxrA) were identified in the IPS-1, BLZ2, and M.Nitro MAGs, and a nitrate reductase closely related to NarG proteins was identified in the BLZ2 MAG. Of the *Methanoperedenaceae* MAGs, only the M.Nitro and BLZ2 MAGs possess a putative nitrite reductase (NrfA) for DNRA. The M.Ferri MAG encodes an assimilatory nitrate reductase (NarB/NasA) most similar to a protein encoded by the *Magnetobacterium casensis* (**Figure 3**). However, in the absence of an annotated nitrite reductase in the M.Ferri MAG, the potential of this microorganism for assimilatory nitrate reduction is unclear.

Multiple MAGs (ASW-2,3,5,6, and Mn-2) were also found to encode putative selenate reductases (SrdA; **Figure 3**), suggesting their ability for Se(VI)-dependent AOM. Recently, a bioreactor enrichment of a member of the genus “*Ca*. Methanoperedens” exhibited AOM activity when nitrate was substituted with selenate (26). However, as no meta-omic analyses was conducted for the community, it is unclear if the dominant “*Ca*. Methanoperedens” possessed a putative selenate reductase, or if it was directly responsible for the observed selenate reduction.

The ASW-1 and ASW-3 MAGs encode a putative sulfur reductase (SreABC). This annotation is supported by its phylogenetic clustering of the catalytic sub-unit with SreA from *Aquifex aeolicus* (**Figure 3**), which has been shown to reduce elemental sulfur, as well as tetrathionate and polysulfide (66). This is the first genomic evidence suggesting that members of the *Methanoperedenaceae* may be involved in respiratory sulfur-dependent AOM and warrants further investigation. ANME-1 have been proposed to couple AOM to the reduction of polysulfide in a biogenic hydrocarbon seep sediment, but this was based on the annotation and high expression of a putative sulfide: quinone oxidoreductase (SQR)(67). Genes for dissimilatory sulfate reduction pathways were absent in the *Methanoperedenaceae* MAGs, consistent with other ANME lineages (68). MGW-1 was recently speculated to directly couple AOM to sulfate reduction utilising assimilatory sulfate reduction pathways. This hypothesis was based on the lack of large MHCs or identifiable alternate electron acceptor complexes encoded in the MAG (29). Several of the *Methanoperedenaceae* MAGs, and those of other ANME lineages, contain candidate genes associated with assimilatory sulfate reduction, but a dissimilatory role for these has not been shown (68).

The M.Nitro MAG encodes two putative reductases belonging to the arsenate reductase (ArrA) and arsenite oxidase (ArxA) group (**Figure 3**). The BLZ2, ASW-1, ASW-4, IPS-1 MAGs also encode reductases that cluster with the M.Nitro ArxA-like sequence. The ArxA protein has been found to be capable of both arsenite oxidation and arsenate reduction (69), which would allow the *Methanoperedenaceae* possessing these ArxA-like proteins to utilise arsenate as a terminal electron acceptor. Proteins encoded by the ASW-3 and “*Candidatus* Acetothermum autotrophicum” (70) (**Figure 3**) form a deep branching clade adjacent to the ArxA and ArrA groups, suggesting these species might also have the potential to respire on arsenic compounds. It is noteworthy that the ASW-1, 3, and 4 MAGs were recovered from a Bangladesh arsenic contaminated groundwater sample (**Table 1**), indicating a role for LGT in their niche-specific adaptation. The possibility of AOM coupled to arsenate (As(V)) reduction has important environmental implications given the wide distribution of arsenic in nature, including subsurface drinking water aquifers (71), and the toxicity and mobility of its reduced form, arsenite (As(III)) (72) (73). Arsenic reduction and mobilisation has been linked to an inflow of organic carbon in contaminated aquifers where methane (~1mM) and arsenate co-occur (74, 75).

Additional putative oxidoreductases clades that are not closely associated with any well characterised CISM proteins were also found in the *Methanoperedenaceae* MAGs. This includes two proteins encoded by the ASW-3 and ASW-6 MAGs that cluster with a protein of unknown function from a *Brocadiales* MAG (76), and the CMD-1 protein that clusters with a protein from *Brocadia fulgida*, an ammonium oxidising and nitrite reducing microorganism (77). In general, given the large range of substrates utilized by the CISM superfamily and the few biochemically characterized proteins, the predicted function of all those annotated in the *Methanoperedenaceae* require empirical verification. Nonetheless, the range of putative CISM superfamily proteins encoded by members of the family likely indicates diverse respiratory strategies that remain to be characterised.

### The diversity of the MHCs in the Methanoperedenaceae

Members of the *Methanoperedenaceae* possess a diverse repertoire of MHCs which have been suggested to facilitate the transfer of electrons from the re-oxidation of MK to metal oxides (9, 10, 78) or direct interspecies electron transfer (DIET) to a syntrophic partner. Analyses of the *Methanoperedenaceae* revealed that they possess between three (MGW-1) and 49 (IPS-1) MHCs (containing at least three CXXCH motifs) with an average of 26 – the highest average of any archaeal family (**Dataset S1F and S1G**). Notably, relatively high numbers of MHCs per genome are almost exclusively found in microorganisms associated with DIET, metal and/or sulfur reduction, such as the *Geobacteraceae* (79) (≤ 87 MHCs)*, Shewanellaceae* (80) (≤ 63 MHCs), *Desulfurivibrionaceae* (20), *Desulfuromonadaceae* (20) and *Defferisomataceae* (81) (≤ 50 MHCs; **Dataset S1G**). Interestingly, seven of the 16 members of the *Methanoperedenaceae* encode MHCs with more than 50 heme binding sites (ASW-5, ASW-6, BLZ2, HGW-1, M. ferri, Mn-1 and Mn-2), with the 113 heme MHC encoded by Mn-2 the largest identified in any microorganism (**Dataset S1F**).

The 414 putative MHCs identified in the *Methanoperedenaceae* MAGs clustered into 82 orthologous protein families (**Figure S8**). Only one protein family (OG0000252) included at least one MHC from each member, which suggests low conservation of these genes within the *Methanoperedenaceae*. Out of the 82 MHC protein families, 14 were identified in at least eight of the 16 MAGs, with five of these found within the conserved MK:cytochrome *c* oxidoreductase clusters. A lack of conservation of MHCs is also observed for the anaerobic metal-respiring genus *Geobacter*, where 14% of the MHCs encoded in six analysed genomes were found to be conserved (61). Thirty-nine of the 82 MHC protein families had significant hits (1e-20, ≥50% AAI) to homologs from diverse lineages across the bacterial and archaeal domains in the GTDB89 database, indicating potential LGT of these genes (**Figure S9**). These lineages notably included the metal reducing *Geobacteraceae* and *Shewanellaceae*, along with the alkane oxidising *Archaeoglobaceae*, Methylomirabilota (NC10), and other ANME-lineages (**Figure S9**).

### Putative function of MHCs in the Methanoperedenaceae

Very few of the *Methanoperedenaceae* MHCs could be associated with a specific function. Two orthologous groups were annotated as nitrite: ammonium oxidoreductases (NrfA) with homologs identified in bacterial MAGs classified to the Anaerolineales (OG0004545; ≥66.3% AAI) and the candidate phylum UBP4 (OG0012490, 64.56% AAI). Several MHCs were also identified as part of the MK:cytochrome *c* oxidoreductase clusters, with homologs observed in members of the archaeal family *Archaeaglobaceae* (OG001557, OG000137, OG0001550, ≥57.3% AAI; **Figure S9**). MHC/S-layer fusion proteins were suggested to mediate the transfer of electrons across the S-layer for marine ANME-2 (ref. 7) and were relatively highly expressed by ‘*Ca*. M. manganicus’ and ‘*Ca*. M. manganireducens’ during AOM coupled to Mn(IV) reduction (12). Conversely, only low expression of MHC/S-layer protein genes encoded by ‘*Ca*. M. ferrireducens’ was observed during AOM coupled to Fe(III) reduction (9). In addition, despite all the *Methanoperedenaceae* MAGs containing S-layer proteins, five do not encode MHC proteins with an S-layer domain (ASW-3, CMD-1, CMD-2, HGW-1 and MGW-1), indicating alternative mechanisms for electron transfer across the S-layer to extracellular MHCs for these species.

Predicted extracellular MHCs are hypothesized to facilitate the final transfer of electrons from the *Methanoperedenaceae* to metal oxides (9). Interestingly, ‘*Ca*. M. manganicus’ and ‘*Ca*. M. manganireducens’ showed differential expression patterns in the complement of shared extracellular MHCs during AOM coupled to Mn(IV) reduction. In addition, no orthologs for the two MHCs highly transcribed by ‘*Ca*. M. ferrireducens’ during AOM coupled to Fe(III) reduction (9) were identified in other members of the *Methanoperedenaceae* (OG0011636 and OG0003254; **Figure S8**), suggesting that BLZ2 utilises a different MHC for iron reduction linked to AOM (10). These observations suggest that the *Methanoperedenaceae* can utilise multiple mechanisms for the reduction of similar metal oxides. Differential expression of conserved MHCs linked to extracellular electron transfer was also observed for different *Geobacteraceae* species enriched on electrodes when exposed to the same surface redox potential (82). As suggested for members of the *Geobacteraceae*, the large MHC repertoire possessed by the *Methanoperedenaceae* may enable adaptation to the use of a range of terminal electron acceptors.

This study has substantially improved the genome coverage of the *Methanoperedenaceae*. Comparative genomic analysis of this lineage highlights a metabolic plasticity not found in other ANME clades. The subsequent ability of members of the family to adapt to the use of terminal electron acceptors across a range of redox potentials likely contributes to their success in diverse environments (**Table 1**). Notably, based on the genome tree (**Figure 1**), and the lack of conservation of MHCs (**Figure S8**), the acquisition of these genes is not congruent with the genome-based phylogeny of the family, suggesting niche specific adaptations as the main driver for these LGT events. While further studies are necessary to verify the general physiology and energy conservation mechanisms of the *Methanoperedenaceae* in different environments, this study provides genomic evidence that members of the family may play key roles in coupling cycling of carbon with selenate, sulfur, and arsenic in addition to nitrogen and metal oxides. Continued sequencing and characterisation of this lineage will reveal the full extent of their metabolic versatility and influence on global biogeochemical cycles.

## Materials and Methods

### Recovery of the genomes from SRA

The NCBI sequence read archive (SRA (83)) was accessed on the 22^nd^ of March 2017 and 14516 datasets classified as environmental metagenomes were downloaded. The metagenomic datasets were screened using GraftM (32) to search for 16S rRNA and *mcrA* gene sequences similar to those from members of the *Methanoperedenaceae*. For datasets where members of the family were detected, all paired-end read sets were trimmed and quality filtered using PEAT v1.2.4 (ref. 84). For genomes, CMD-1 and CMD-2, SRR5161805 and SRR5161795 reads were coassembled using Metaspades, version 3.10.0 using the default parameters (85). For the ASW genomes, SRR1563167, SRR1564103, SRR1573565, and SRR1573578 reads were coassembled using Metaspades, version 3.10.0 with default parameters (85). Mapping of quality reads was performed using BamM v1.7.3 with default parameters (https://github.com/Ecogenomics/BamM). Metagenomic assembled genomes were recovered from the assembled metagenomes using uniteM v0.0.14 (https://github.com/dparks1134/UniteM). The *Methanoperedenaceae* MAGs were further refined by reassembling the mapped quality trimmed reads with SPAdes using the –careful and –trusted-contigs setting. Additional scaffolding and resolving ambiguous bases of the MAGs was performed using the ‘roundup’ mode of FinishM v0.0.7 (https://github.com/wwood/finishm). Completeness and contamination rates of the population bins were assessed using CheckM v1.0.11 (ref. 33) with the ‘lineage wf’ command. The genomes assembled in this study have been deposited in NCBI under the accession numbers SAMN10961276-SAMN10961283.

### Functional annotation

For all MAGs, open reading frames (ORF) were called and annotated using Prokka v.1.12 (ref. 86). Additional annotation was performed using the blastp ‘verysensitive’ setting in Diamond v0.9.18 (https://github.com/bbuchfink/diamond.git) against UniRef100 (accessed September 2017) (87), clusters of orthologous groups (COG) (88), Pfam 31 (ref. 89) and TIGRfam (Release: January 2014) (90). ORFs were also diamond blastp searched against Uniref100 (accessed September 2017) containing proteins with KO ID. The top hit for each gene with an e-value <1e^-3^ was mapped to the KO database (91) using the Uniprot ID mapping files. Genes of interest were further verified using NCBI’s conserved domain search to identify conserved motif(s) present within the gene (92). Psortb v3.0 (ref. 93) was used to predict subcellular localisation of the putative proteins. Pred-Tat was used to predict putative signal peptides (94). Putative multi-heme *c*-type cytochromes (MHCs) were identified by ORFs possessing ≥3 CXXCH motifs. Putative MHCs were subsequently searched for cytochrome *c*-type protein domains using hmmsearch (HMMER v.3.1) (95) with PfamA (96).

### Construction of genome trees

The archaeal genome tree was constructed using GTDB-Tk (GTDBtk v0.2.2, https://github.com/Ecogenomics/GTDBTk/releases) with a concatenated set of 122 archaeal-specific conserved marker genes inferred from genomes available in NCBI (NCBI RefSeq release 83) (34). Marker genes were identified and aligned in each genome using HMMER v.3.1 (ref. 95), concatenated, and trees were constructed using FastTree V.2.1.8 (ref. 97) with the WAG+GAMMA models. Support values were determined using 100 nonparametric bootstrapping with GenomeTreeTK. The trees were visualised using ARB (98) and formatted using Adobe Illustrator (Adobe, USA).

### Construction of 16S rRNA gene tree

The 16S rRNA gene was identified in MAGs and used to infer taxonomic assignment of the population genome implementing the SILVA 16S rRNA gene database (Version 132). Sequences were aligned with 426 16S rRNA gene sequences retrieved from the SILVA database using SSU-align v0.1 (ref. 99). The phylogenetic tree was constructed using FastTree V2.1.8 (ref. 97) with the Generalised Time-Reversible and GAMMA model. Support values were determined using 100 nonparametric bootstrapping. The trees were visualised using ARB (98) and formatted using Adobe Illustrator.

### Calculation of amino acid identity

The *Methanoperedenaceae* MAGs identified in this study were compared to publicly available genomes of the family. Average amino acid identity (AAI) between the genomes was calculated using orthologous genes identified through reciprocal best BLAST hits using compareM v0.0.5 (https://github.com/dparks1134/CompareM).

### Identification of orthologous proteins

Homologous proteins across all the *Methanoperdenaceae*, GoM-Arc I, ANME-2a, ANME-2c MAGs were identified with OrthoFinder (100) v2.3.3 using default parameters. Gene counts of orthologous groups containing MHCs were used as input for a heatmap using the pheatmap package in R and hierarchical clustering was performed using ward.D2 (ref. 101).

### Construction of gene trees

Genes of interest in the *Methanoperedenaceae* MAGs were compared against proteins from GTDB v83 database (34) using the genetreetk ‘blast’ command to identify closely related sequences. For the generation of the gene tree for catalytic subunits of the CISM superfamily, curated protein sequences were also added in the analysis. Accession numbers and amino acid sequences are included in **Dataset S1B**. For the generation of the gene tree for the catalytic subunits of the Group 1 and Group 3 NiFe dehydrogenase, curated sequences from Greening *et al*., (102) were included in the analysis. Accession numbers and amino acid sequences can be found in **Dataset S1C**. The sequences were subsequently aligned using mafft v7.221 (ref. 103) with the –auto function and the alignment trimmed using trimal v1.2 (https://github.com/scapella/trimal) ‘-automated1’ option. A phylogenetic tree was constructed using RAxML v8.2.9 (ref. 104) with the following parameters: raxmlHPC-PTHREADS-SSE3 -T 30 -m PROTGAMMALG -p 12345. Bootstrap values were calculated via non-parametric bootstrapping with 100 replicates. The trees were visualised using ARB (98) or iToL (105) and formatted using Adobe Illustrator (Adobe, USA).

### Network analysis of MHCs

Putative multi-heme *c*-type cytochromes (MHCs) from GTDB v89 database were identified by ORFs possessing ≥3 CXXCH. Putative MHCs were subsequently searched for cytochrome *c*-type protein domains using hmmsearch (HMMER v.3.1) (95) with PfamA (96). Proteins from each *Methanoperedenaceae* orthogroup were blasted against the GTDB v89 MHC protein database using DIAMOND with an evalue cutoff of 1e-20 and ≥50% AAI. The result was visualised in Cytoscape v3.7.1, removing clusters that contained only, or no, *Methanoperedenaceae* homologs.

## Acknowledgements

This work was supported by the Australian Research Council (ARC) (FT170100070) and the U.S. Department of Energy’s Office of Biological Environmental Research (DE-SC0016469). AOL was supported by an ARC Australian Postgraduate Award. We thank the AWMC team, particularly Shihu Hu and Zhiguo Yuan, for their ongoing collaboration working on various “*Ca*. Methanoperedens” enrichments.

## Competing interests

The authors have nothing to disclose.

## Supplemental material legends

**Figure S1. Average amino acid identity (AAI%) for the *Methanoperedenaceae* genomes.** AAI was calculated between each pair of genomes using CompareM.

**Figure S2. 16S rRNA gene based phylogenetic placement of the *Methanoperedenaceae* MAGs.** The 16S rRNA genes extracted from the *Methanoperedenaceae* MAGs from this study are highlighted in red. Support values calculated via non-parametric bootstrapping. The scale bar represents changes per nucleotide position.

**Figure S3. Phylogenetic analysis of methyl-coenzyme reductase subunit A (McrA).** Putative genes recovered from the *Methanoperedenaceae* are highlighted in red. The gene tree was inferred using maximum likelihood and support values calculated via non-parametric bootstrapping. The scale bar represents amino acid changes.

**Figure S4. Phylogenetic analysis of the subunits of the NiFe hydrogenases annotated in the *Methanoperedenaceae* genomes. A.** Analysis of the catalytic subunits of the energy-converting NiFe hydrogenases. **B.** Analysis of the *b*-type cytochrome in the Group 1 NiFe hydrogenases. Putative genes recovered from the *Methanoperedenaceae* are highlighted in red. The gene trees were inferred using maximum likelihood and support values calculated via non-parametric bootstrapping. The reference sequences of Group 1 and Group 3 NiFe hydrogenases were acquired from Greening *et al*., (C. Greening, A. Biswas, C. R. Carere, C. J. Jackson, M. C. Taylor, M. B. Stott, G. M. Cook and S. E. Morales, ISME J 10: 761-777, 2016, https://doi.org/10.1038/ismej.2015.153) and the GTDB v83 reference sequences (D.H. Parks, M. Chuvochina, D. W. Waite, C. Rinke, A. Skarshewski, P.-A. Chaumeil and P. Hugenholtz, Nat Biotechnol 36: 996-1004, 2018, https://doi.org/10.1038/nbt.4229). The scale bars represent amino acid changes.

**Figure S5. Subunit compositions of the Fpo dehydrogenase protein complexes and theoretical bioenergetics of energy metabolism in ANME-2a and *Methanoperedenaceae*. A.** Fpo subunit components for the ANME-2a and ASW-3 genomes (top left) and the other members of the *Methanoperedenaceae* (bottom left). The utilization of different electron carriers shows greater biochemical energetic gains based on more potential proton translocation. The colours orange and green depict Methanosarcinales-like and non-Methanosarcinales-like subunits. **B.** Theoretical redox potential drop when utilizing MP (left) or MK (right) during F420H2 and Fd^2-^ oxidation. This is due to differences between the membrane-bound electron carriers’ redox midpoint potential (Em) of −80mV and −165mV for MK and MP, respectively (M., Tietze, A. Beuchle, I. Lamla, N. Orth, M. Dehler, G. Greiner and U. Beifuss, Chembiochem 4: 333-335, 2003, https://doi.org/10.1002/cbic.200390053; Q.H. Tran and G. Unden, Eur. J. of Biochem. 251: 538-543, 1998, https://doi.org/10.1046/j.1432-1327.1998.2510538.x).

**Figure S6. Phylogenetic analysis of the Fpo subunits annotated in the *Methanoperedenaceae* genomes. A.** FpoA **B.** FpoB **C.** FpoC **D.** FpoD **E.** FpoH **F.** FpoI **G.** FpoJ1 **H.** FpoJ2 **I.** FpoK **J.** FpoL **K.** FpoM **L.** FpoN **M.** FpoO. Putative genes recovered from the *Methanoperedenaceae* are highlighted in red. The gene trees were inferred using maximum likelihood and support values calculated via non-parametric bootstrapping. Reference genes and the taxonomy are from the GTDB v83 database (D.H. Parks, M. Chuvochina, D. W. Waite, C. Rinke, A. Skarshewski, P.-A. Chaumeil and P. Hugenholtz, Nat Biotechnol 36: 996-1004, 2018, https://doi.org/10.1038/nbt.4229).

**Figure S7. Phylogenetic analysis of the subunits of the MK:cytochrome oxidoreductases annotated in the *Methanoperedenaceae* MAGs. A.** Analysis of the NrfD subunits. **B.** Analysis of the b-type cytochromes. Bootstrap values for the maximum-likelihood trees were determined using non-parametric bootstrapping with 100 replicates. The scale bars represent amino acid changes.

**Figure S8. Abundance profiles for the MHC orthologous protein families annotated in the *Methanoperedenaceae* MAGs.**

**Figure S9. Network analysis of MHC orthologous protein families in *Methanoperedenaceae*.** Each cluster represents related MHCs. The colour of the nodes represents the taxonomic lineage based on GTDB classification. The size of the nodes represents the number of CXXCH heme binding motifs identified in the proteins. The thickness of the lines represents amino acid identity between the two nodes. The shaded boxes represent the orthologous protein families.

**Dataset S1. Sequences, identifiers and statistics for genes used in the comparative analyses of the *Methanoperedenaceae* MAGs. A.** Genes encoding proteins involved in the methane oxidation pathway, energy conservation, and other metabolic pathways as shown in Figure 2. **B.** Amino acid sequences used in the CISM superfamily gene tree (Figure 3). Amino acid sequences include curated sequences from Swiss-Prot and Castelle *et al*., (C.J. Castelle, L. A. Hug, K. C. Wrighton, B. C. Thomas, K. H. Williams, D. Wu, S. G. Tringe, S. W. Singer, J. A. Eisen and J. F. Banfield, Nat commun 4: 2120, 2013, https://doi.org/10.1038/ncomms3120) and closely related sequences from GTDB r83 protein reference database (D.H. Parks, M. Chuvochina, D. W. Waite, C. Rinke, A. Skarshewski, P.-A. Chaumeil and P. Hugenholtz, Nat Biotechnol 36: 996-1004, 2018, https://doi.org/10.1038/nbt.4229). **C.** Amino acid sequences used in the catalytic subunits of the energy-converting NiFe hydrogenase. Amino acid sequences include curated sequences from Greening *et al*., (C. Greening, A. Biswas, C. R. Carere, C. J. Jackson, M. C. Taylor, M. B. Stott, G. M. Cook and S. E. Morales, ISME J 10: 761-777, 2016, https://doi.org/10.1038/ismej.2015.153) and closely related sequences from GTDB r83 protein reference database. **D.** Genes encoding putative NiFe hydrogenase maturation proteins. **E.** Best blastp hits of Fpo dehydrogenase subunits to the IMG database. Blastp hits shows divergent Fpo subunits are present in the *Methanoperedeneceae* MAGs as seen in Figure S6. Top blast hits to Methanoperedens-like protein sequences were excluded. **F.** General statistic of multi-heme *c*-type cytochromes (MHCs) in the ANME genomes. **G.** MHC general statistics for all bacterial and archaeal families in the GTDB v89 database.

